# Nicotinamide promotes differentiation of pancreatic endocrine progenitors from human pluripotent stem cells through poly (ADP-ribose) polymerase inhibition

**DOI:** 10.1101/2020.04.21.052951

**Authors:** Curtis Woodford, Ting Yin, Huntley H Chang, Romario Regeenes, Ravi N Vellanki, Haneesha Mohan, Bradly G Wouters, Jonathan V Rocheleau, Michael B Wheeler, Peter W Zandstra

## Abstract

The generation of pancreatic endocrine progenitor cells is an important step in the differentiation of beta cells from human pluripotent stem cells (hPSC). This stage is marked by the expression of Nkx6.1, a transcription factor with well understood downstream targets but with unclear upstream regulators. In hPSC differentiation, Nkx6.1 is strongly induced by nicotinamide, a derivative of vitamin B3, which has three known functions within a cell. Nicotinamide inhibits two classes of enzymes known as poly-ADP-ribose polymerases (PARPs) and sirtuins. It also contributes to the cellular pool of nicotinamide adenosine deoxynucleotide (NAD^+^) after conversion in the nicotinamide salvage pathway. Induction of Nkx6.1 expression in pancreatic endocrine progenitors by nicotinamide was mimicked by 3 PARP inhibitors (PJ34, olaparib, NU1025). Small molecule inhibition of the nicotinamide salvage pathway reduced Nkx6.1 expression but not Pdx1 expression and caused alteration in NAD^+^/NADH ratio. Nkx6.1 expression was not affected by sirtuin inhibition. Metabolic profiling of differentiating pancreatic and endocrine progenitors showed that oxygen consumption increases as differentiation progresses, and that nicotinamide reduces oxygen consumption rate. Expression of Nkx6.1 and other beta cell related genes, including Ins2 and Pdx1 increased in mouse islets after exposure to nicotinamide. In summary, nicotinamide induced Nkx6.1 expression in differentiating human pancreatic endocrine progenitors through inhibition of the PARP family of enzymes. Nicotinamide administration was also associated with increased NAD+/NADH ratio, without affecting Nkx6.1 expression. Similarly, the association between nicotinamide and Nkx6.1 expression was also seen in isolated mouse islets. These observations show a link between the regulation of beta cell identity and the effectors of NAD^+^ metabolism, suggesting possible therapeutic targets in the field of diabetes.

## Introduction

Pancreatic progenitors derived from human pluripotent stem cells (hPSC) offer unique insight into pancreatic beta cell development and also offer a cell replacement strategy for treating type 1 diabetes [1],[2]. Multiple differentiation protocols have been established for deriving pancreatic progenitors from hPSC [3],[4]. One of these uniquely uses high dose nicotinamide to induce Nkx6.1 expression during the conversion from pancreatic to endocrine progenitor [5]. Nicotinamide is a derivative of vitamin B3 and has been shown to have multiple roles in the cell. It is converted to NAD^+^ in the nicotinamide salvage pathway, which is then used in many ways, including being consumed by the mitochondria, sirtuins and PARP family of enzymes [6]. The sirtuin and PARP families of enzymes use NAD^+^, but are also inhibited by nicotinamide, which leads to both activating and inhibiting effects depending on the activity level of the nicotinamide salvage pathway [7],[8].

The sirtuins are a seven-member family of enzymes with different subcellular localizations such as the nucleus or mitochondria involved in gene and metabolic regulatory pathways [9]. Sirtuin inhibition has been associated with vascular disease, diabetes, and metabolic syndrome [10]. Their activation is thought to extend lifespan, increase oxidative phosphorylation, and improve insulin responsiveness [11]. The PARP family of enzymes consists of 17 members, of which the best studied are PARP1, 2, and 3 [12]. They were originally discovered as DNA repair enzymes, but have recently been found to poly-ADP ribosylate thousands of proteins and DNA sequences in the cell [13]. They have functions in gene expression, metabolism, apoptosis, and DNA repair processes. PARP inhibitors were originally found to be protective against streptozotocin-induced diabetes, suggesting a role in coping with cellular metabolic stress [14].

Nkx6.1 is a transcription factor expressed in the beta-cells of the islet, and appears to have function closely linked to the energy status of the cell [15]. For instance, Nkx6.1 knockout reduces glycolytic flux and completely abolishes insulin expression [16]. It is also one of the first transcription factors to be downregulated in the setting of type 2 diabetes in both mice and humans [17]. Despite this central role in beta cell function and dysfunction, regulation of Nkx6.1 expression is not well understood although downstream targets of this transcription factor have been well identified [18].

Here we show that Nkx6.1 is controlled by PARP inhibition during human endocrine pancreatic differentiation of hPSC. We show that Nkx6.1 expression during hPSC differentiation is not dependent on sirtuin inhibition, and that alterations to oxidative metabolism occur as cells become more mature pancreatic endocrine progenitors. These changes are similar to those that occur to isolated mouse islets, suggesting that the effects of nicotinamide are preserved across species. The results in this paper suggest a linkage between the PARP family of enzymes and beta cell dedifferentiation in type 2 diabetes.

## Methods

### hPSC culture and differentiation

hESC were expanded on mouse embryonic feeder seeded, 0.2% gelatin (Sigma) coated 6 well plates in media containing 80% DMEM/F12 (Gibco), 20% knockout serum replacement (Gibco), 1% non-essential amino acids, 0.5% (which is 5,000 units/mL of penicillin and 5,000 μg/mL of streptomycin (0.5% Gibco), and 1% GlutaMAX supplement (Gibco), 0.1mM 2-mercaptoethanol (Sigma) and 10 ng/mL bFGF (Pepro Tech). They were split at a 1:10-1:20 ratio. Prior to differentiation they were seeded into 6 well or 12 well plates treated with Geltrex (Gibco) diluted 1:50 with DMEM/F-12. They were passaged at 90% confluency at a 1:6 ratio. Differentiation was initiated at 90% confluence, usually after 72 hours of culture on Geltrex (Gibco) coated plates in Nutristem hPSC XF media (Biological Industries).

Endoderm base media consisted of RPMI 1640 with added 1% glutamax and 1% penicillin-streptomycin. On day 1, CHIR 99021 (Reagent Direct) was added at 2 μM with activin A made in house at a 5 μL/mL ratio. This was approximated using commercially available activin A to be 100 ng/mL concentration. Day 2 and 3 media consisted of endoderm differentiation (Sigma) base media with added activin A (5 μL/mL), bFGF at 5 μg/mL and ascorbic acid 50 ug/ml. Day 4 and 5 media consisted of pancreatic progenitor base media composed of DMEM, 1% B27 (Gibco), 1% glutamax and 1% penicillin-streptomycin. To this was added FGF10 (R & D Systems) at 50 ng/mL for days 4 and 5. On day 6 and 7, FGF10 (50 ng/mL), retinoic acid (2 μM, Sigma), LDN192168 (2 nM, Reagent Direct), KAAD-cyclopamine (2.5 μM, Toronto research chemicals), PDBu (4 μM, Sigma) were added. On days 8 and 10, pancreatic progenitor base media plus 50 ng/mL EGF (R & D Systems), 2 nM LDN192168, and either 0 or 10 mM nicotinamide (Sigma) were added. Olaparib (Sigma) was added at 10 nM, PJ34 (Sigma) at 4 μM, and Nu1025 (Tocris) at 10 μM.

### Flow cytometry

Cells were dissociated with TrypLE for 5 min, then washed with an equal volume of 2% FBS in HBSS (referred to as HF). They were centrifuged and washed with a second volume of HF. CXCR4 and ckit conjugated antibodies (BD Biosciences) were added to the cells in HF at 1:100 and 1:50 ratios, respectively. Cells were incubated for 20 min at room temperature, then washed with 1 mL of HF and fluorescence intensity was measured using a BD FACS Canto flow cytometer. If performing intracellular staining, cells were suspended in 4% paraformaldehyde in PBS, 100 uL for 20 min at room temperature. They were washed with 5% FBS in PBS and resuspended in methanol kept on ice for 2 min. They were then washed with 5% FBS in PBS and resuspended in the same solution for blocking at 4 C. Primary antibodies Nkx6.1 (1:100, DSHB) and Pdx1 (1:100, R&D Systems) were added and incubated for 30 minutes. Cells were washed with 5% FBS in PBS and strained for 30 minutes at room temperature with secondary antibodies (Molecular Probes). They were then washed twice and fluorescence intensity was measured on BD FACS Canto.

### NAD^+^/NADH ratio measurement

hPSC were differentiated to the end of Stage 4, with 4 μM PJ34, 10 μM nicotinamide, or 1 nM FK866 with water or DMSO added in equal volume to the test condition as a negative control. Cells were dissociated with TrypLE at 4°C for 5 min and pelleted after 2x dilution with PBS + 5% FBS at 4°C. Cells were kept on ice for the remainer of the assay. The EnzyChrom ™ NAD^+^/NADH Assay Kit (BioAssay Systems E2ND-100) was used to quantify the NAD^+^/NADH ratio using absorption of 565 nm light after a lactate dehydrogenase cycling reaction involving formazan (MTT) as a reducing agent, according to the manufacturer’s instructions.

### 2-Photon NAD(P)H Imaging

Cells were differentiated to day 9 according to the differentiation protocol above. They were dissociated with TrypLE and re-plated on Matrigel-coated 35 mm Mattek glass bottom dishes at a density of 500 000 cells per dish. Cells were incubated in stage 4 media without nicotinamide overnight. Cells were imaged using a Zeiss LSM710 confocal equipped with a Ti:Saph two-photon laser tuned to 705 nm at 2.5 mW, a 40x/1.3 NA oil immersion lens, and an infrared-blocked band pass filter (385-550 nm). Images consisted of 512×512 pixels sized at 0.14 μm/pixel and were taken with a pixel dwell time of 6.30 μs. The cells were incubated for 10 min with 10 μM nicotinamide, 4 μM PJ34, FCCP, or rotenone and NAD(P)H autofluorescence measurements were repeated. A ratio of the relative fluorescence units prior to reagent and after reagent addition was calculated.

### Western Blotting

Cells at day 12 of differentiation were lysed with cell lysis solution containing fresh protease inhibitor at 4 C. Lysate was spun down at 14000 rpm and protein concentration was determined using the Bradford assay as detailed in the manufacturer’s kit (Biorad). Samples were denatured with SDS and methylene blue, and 30 mg of protein were added to each lane of the premade gel. Electrotransfer was performed using the iBlot 2 (Life Technologies) system according to manufacturer’s instructions. Digitization and development of the western blot was performed chemiluminescence using a digitizer. Image analysis of band intensity was performed using Image J.

### Quantitative Real Time Polymerase Chain Reaction

Cells were dissociated with TrypLE and centrifuged, supernatant was aspirated. RNA was isolated using PureLink RNA mini kit (Life Technologies) according to manufacturer’s instructions. Briefly, cells were centrifuged and supernatant was discarded. Cells were lysed in the presence of B-mercaptoethanol, and RNA was isolated using a column extraction method. RNA content was measured in RNase free distilled water, and cDNA was made from this sample using a reverse transcription reaction. qRT-PCR was performed for each transcript using SYBR Green (Life Technologies) as the detection method.

### Seahorse metabolic profiling

Cells were differentiated to either day 9 or day 12 according to the differentiation protocol above. They were dissociated with TrypLE and re-plated at a density of 40 000 cells per well in a Matrigel-coated (BD Biosciences) Seahorse assay plate (Agilent). Cells were incubated in stage 4 media overnight. Cells were exposed sequentially to palmitate, oligomycin, FCCP, and antimycin, or oligomycin, FCCP, and antimycin in basal media containing no bicarbonate buffer. Fluorescence intensity from oxygen and pH sensitive fluorophores in each well are converted to oxygen consumption and acid production rates under each condition. At the conclusion of the assay, DNA content was measured using an intercalating fluorophore, the intensity of which was correlated with cell number through a standard curve.

### Magnetic sorting

EasySep (Stem Cell Technologies) was used to isolate GP2 positive cells from GP2 negative cells. Briefly, cells were dissociated at day 12 and incubated for 15 min with PE-conjugated GP2 antibody (Novus Biologicals) at a 1:50 ratio. Cells were washed and incubated with anti-PE antibody conjugated to streptavidin. Biotin conjugated magnetic nanoparticles were added and the cells were exposed to a cylindrical magnet. GP2 negative cells were decanted and GP2 positive cells remained in the tube. Each group was either replated for the Seahorse assay or were lysed and used for western blotting. Part of each separation was replated and used for measurement of Nkx6.1 and Pdx1 flow cytometry.

### Animal Care

C57BL/6J male mice were obtained from Charles River at the age of 8 weeks for in vitro studies. After receiving mice were kept one week for acclimatization. Mice were housed in groups of 2-4 at 22°C-24°C using 12 hr light/12 hr dark cycle in Division of Comparative Medicine (DCM) facility, Faculty of Medicine, University of Toronto. Mice were fed a normal chow diet. All mouse procedures were conducted in compliance with protocols approved by the Animal Care Committee at the University of Toronto and the guidelines of the Canadian Council of Animal Care.

### Islet isolation

Eight week old C57BL/6 male mice were anesthetized using isofluorane. Islets were isolated through pancreatic perfusion and collagenase digestion as previously described [34]. Islets were hand picked three times and allowed to recover overnight in RPMI (1640; Sigma) supplemented with 10% FBS and 1% Pen/Strep prior to analysis. They were counted into equal numbers for treatment with either nicotinamide or water. RNA for qRT-PCR was isolated with a micro kit (Life technologies) according to the procedure detailed above.

### Statistical analysis

Unpaired or paired t-tests were performed for statistical analysis of data in figures 1–5, supplementary figure 3. One-way ANOVA was used for supplementary figure 1.

**Figure 1:**
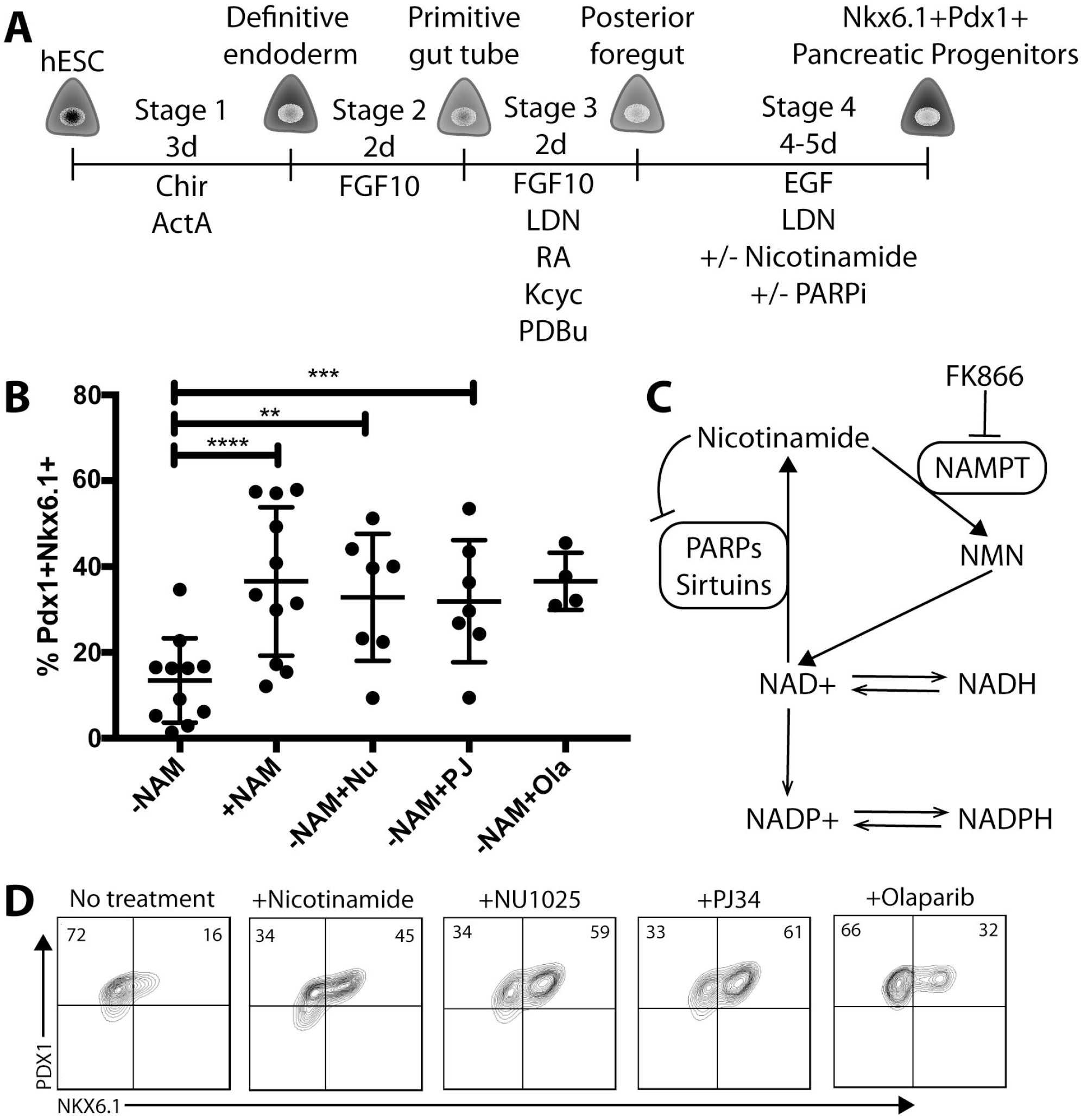
Summary of differentiation protocol and nicotinamide effect on Nkx6.1 expression. A, hPSC are differentiated from pluripotency to pancreatic progenitors over the course of 11-12 days. B, Effects of nicotinamide and PARP inhibitors on Nkx6.1+Pdx1+ expression in pancreatic progenitors at day 12 of differentiation. Dot plot of Pdx1+Nkx6.1+ populations at day 12 measured by flow cytometry after not being treated with nicotinamide (-NAM), treated with nicotinamide (+NAM), or treated with PARP inhibitors Nu1025 (+Nu), PJ34 (+PJ), and olaparib (+Ola). C, A simplified graphic illustrating the nicotinamide salvage pathway, the functions of PARPs and sirtuins, as well as where the small molecule inhibitor FK866 inhibits NAMPT. D, Representative flow cytometry plots of Pdx1 and Nkx6.1 expression with each Parp inhibitor. * indicates p < 0.05, ** indicates p < 0.01, *** indicates p < 0.001, **** indicates p< 0.0001 in a paired Students t-test.

## Results

### Nicotinamide induces Nkx6.1 expression in human pancreatic endocrine progenitors

hPSC were differentiated through an 11 day staged differentiation protocol that has been previously published [5] (Fig. 1A). After 3 days of differentiation with activin A, and induction with 1 day of a GSK3beta inhibitor, >90% of cells co-expressed the surface proteins CXCR4 and ckit (Stage 1). The co-expression of these proteins was found to be highly correlated with the expression of Sox17 and FoxA2, two transcription factors that mark definitive endoderm expression [19]. The cells were then differentiated towards an anterior foregut pathway with FGF10 (Stage 2), and pancreatic differentiation was induced with retinoic acid, BMP and sonic hedgehog inhibition (Stage 3). Endocrine progenitor differentiation was carried out with epidermal growth factor (EGF), BMP inhibition, and nicotinamide (Stage 4). After stage 4, cells were 20-60% double positive for the transcription factors Pdx1 and Nkx6.1, indicating an endocrine progenitor state (Fig. 1B). The absence of nicotinamide decreased the percentage of cells expressing Nkx6.1 by approximately 50%, showing a strong relationship between treatment with nicotinamide and Nkx6.1 induction.

### PARP inhibitors induce Nkx6.1 expression in human pancreatic endocrine progenitors

Nicotinamide serves as a precursor to NAD^+^ via nicotinamide mononucleotide (NMN) in the nicotinamide salvage pathway, and as an inhibitor of the sirtuin and PARP classes of proteins (Fig. 1C). It is unclear which of these 3 effects is predominating when nicotinamide causes Nkx6.1 expression. Multiple PARP inhibitors mimic the effects of nicotinamide on the percentage of Pdx1 and Nkx6.1 double positive cells during pancreatic differentiation (Fig. 1B and 1D). Nicotinamide is a pan-PARP inhibitor, while NU1025, PJ34, and olaparib are derivatives of nicotinamide that have been selected to specifically target PARP1 and PARP2. We found that the PARP1/2 inhibition caused by PJ34 and NU1025 was required for the robust expression of Nkx6.1 during pancreatic endocrine cell differentiation.

### Nicotinamide salvage pathway perturbation affects Nkx6.1 expression levels

Nicotinamide inhibits the PARP family and may also change the NAD^+^/NADH ratio within differentiating cells due to production of NAD^+^ through the nicotinamide salvage pathway (Fig. 1C). The NAD^+^/NADH ratio within a cell is indicative of the redox state, affecting metabolism and ADP-ribosylation, which has impacts on cell signaling, DNA repair, gene regulation, and apoptosis. We inhibited nicotinamide salvage using FK866, an inhibitor of nicotinamide phosphoribosyltransferase (NAMPT). This completely abolished Nkx6.1 expression but maintained Pdx1 expression (Fig. 2A). We hypothesized that PARP inhibition caused by nicotinamide or PJ34 would cause changes in the NAD^+^/NADH ratio which could be reversed by the inhibition of the nicotinamide salvage pathway by FK866. NAD^+^/NADH ratio was measured using a colorimetric assay based on a lactate dehydrogenase cycling reaction, with very little interference from the NADP^+^/NADPH ratio. Nicotinamide significantly increased the NAD^+^/NADH ratio as expected (Fig. 2B). Addition of FK866 in the presence of nicotinamide abolished this change in NAD^+^/NADH ratio (Fig. 2C), most likely due to the reduction in nicotinamide salvage activity through its inhibition of NAMPT. PJ34, a PARP1/2 inhibitor did not significantly change the NAD^+^/NADH ratio (Fig. 2D). It is probable that the effects on the NAD^+^/NADH ratio were more robust with nicotinamide compared to PJ34 because nicotinamide also serves as a source of NAD^+^ whereas PJ34 does not. These observations suggest that 1) the relatively large perturbation to the cellular NAD^+^/NADH ratio represented by the addition of FK866 is detrimental to Nkx6.1 expression but not Pdx1 expression, 2) subtle differences in NAD^+^/NADH ratio represented by PJ34 compared to nicotinamide do not affect Nkx6.1 expression levels, 3) uninhibited functioning of the nicotinamide salvage pathway is necessary for Nkx6.1 expression.

**Figure 2:**
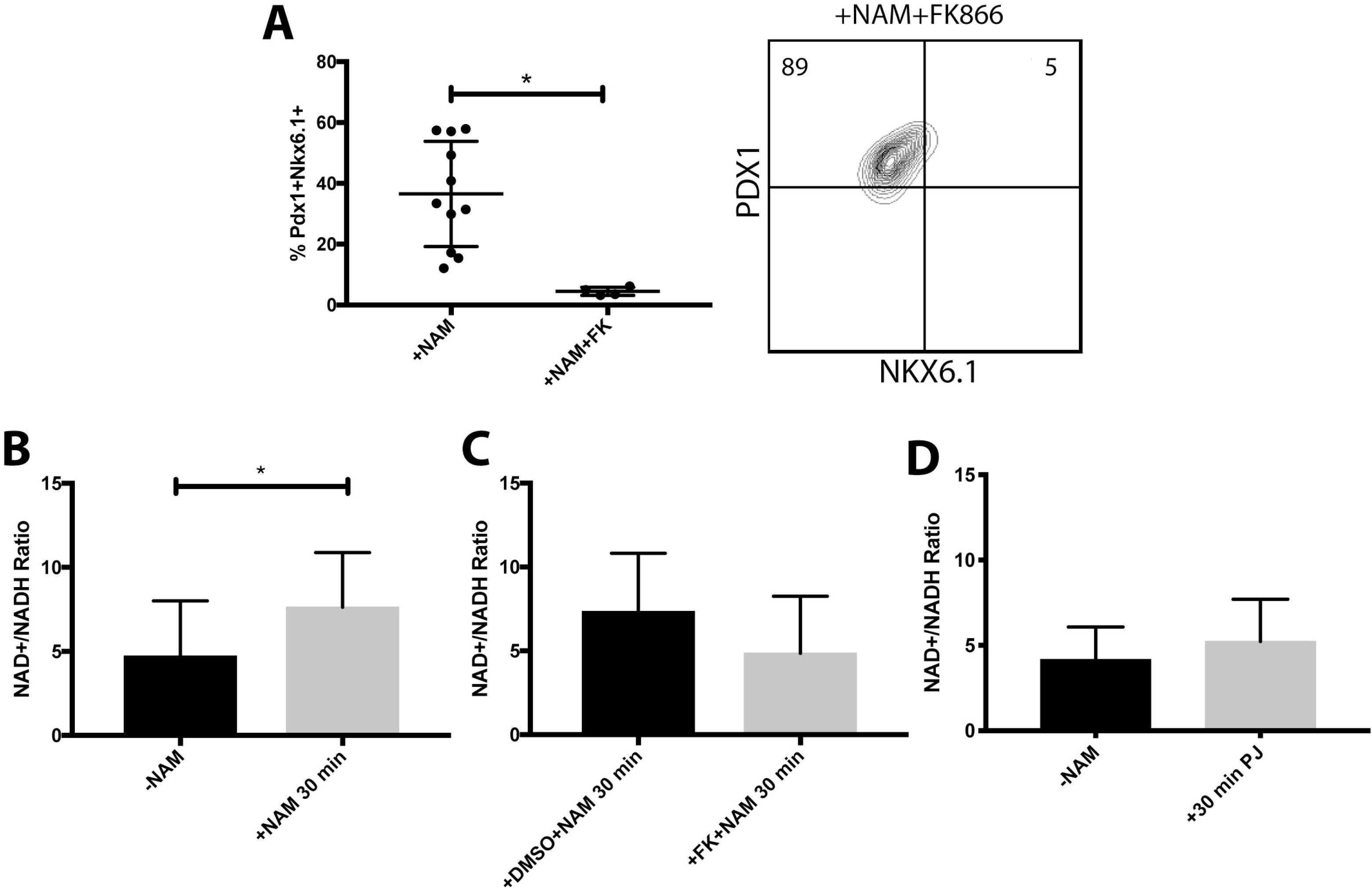
Effects of nicotinamide administration on NAD^+^/NADH ratio. A, FK866 administration with nicotinamide abolishes Nkx6.1 expression. Representative flow cytometry plot of Nkx6.1 expression after FK866 administration. C, NAD^+^/NADH ratio is increased after nicotinamide administration. D, NAD^+^/NADH ratio is unchanged after nicotinamide administration with FK866. E, NAD^+^/NADH ratio is unchanged by PJ34 administration. * indicates p < 0.05 in a paired Students t-test.

Two-photon microscopy allows for subcellular autofluoresence measurement of NAD(P)H levels (NADH + NADPH), which is in dynamic equilibrium with NAD(P)^+^ [20] [21]. We measured the NAD(P)H level in the mitochondria and nucleus of cells differentiated to day 10 (Fig. 1A). Nicotinamide and PJ34 showed no significant effects on the NAD(P)H level in either the nucleus or mitochondria (Suppl. Fig. 1A). The positive control in this experiment was FCCP (Carbonyl cyanide-4-(trifluoromethoxy)phenylhydrazone), a decoupler of mitochondrial oxidative phosphorylation which decreased NAD(P)H levels although not at a statistically significant level (Suppl. Fig. 1B). Rotenone was expected to have the opposite effect but was not observed to statistically significantly increase NAD(P)H levels (Suppl. Fig. 1B and 1C). The lack of robust response to these well characterized molecules suggests that hPSC at this stage of differentiation have immature mitochondrial metabolism, which has been shown in hPSC at earlier stages of development [22], [23].

### Sirtuin inhibition is not necessary for Nkx6.1 expression

Nicotinamide is also an inhibitor of the sirtuin family of enzymes, which use NAD^+^ as a cofactor in lysine deacetylation of many different cellular proteins (Fig. 3A) Human embryonic stem cells were differentiated to day 12 and lysine acetylation levels were measured using Western blotting. Differentiation in the presence of nicotinamide globally increased the amount of acetylated lysine (Fig. 3B and 3C) indicating net-reduced sirtuin family activity. Culture in the presence of PJ34, in contrast, did not have a discernible effect on the amount of acetylated lysine (Fig 3C). EX527 is a targeted inhibitor of sirtuin 1 and 2, and also significantly increased the amount of acetylated lysine in day 12 cells (Fig 3C). We hypothesized that an Nkx6.1 enriched population may have altered sirtuin family activity, so cells exposed to nicotinamide were magnetically sorted into an Nkx6.1 enriched population using the surface antibody GP2 (Suppl. Fig. 2), which has been shown to enrich for endocrine progenitors [24], [25]. Levels of acetylated lysine were not significantly reduced in GP2+ cells compared to GP2-cells, indicating similar levels of sirtuin activity in both populations (Fig. 3C). SRT1720, controversially reported to be an activator of sirtuin 1 [26], did not decrease acetylated lysine levels in this assay (Fig 3C). Given that Nkx6.1 is induced by PJ34 and nicotinamide but not EX527, these data show that that sirtuin inhibition is not necessary for Nkx6.1 expression, and that the endocrine progenitor enriched population has similar sirtuin function to non-endocrine fated cells.

**Figure 3:**
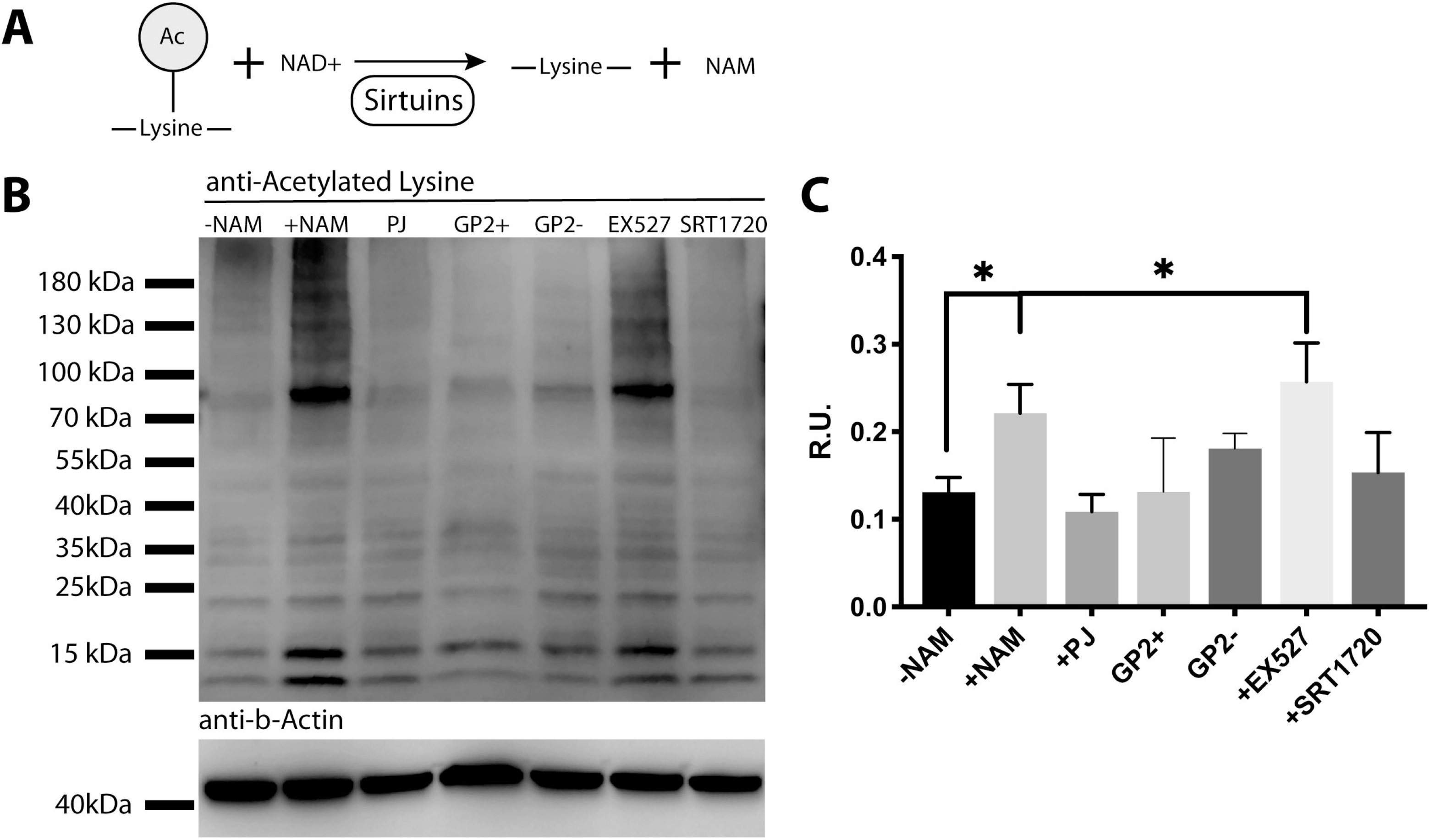
Nicotinamide administration inhibits global lysine deacetylation. A, Sirtuins deacetylate lysine moieties on proteins using NAD^+^ as a cofactor, creating nicotinamide as a product. B, Western blot showing global acetylated lysine levels after adding 30 ug per lane of protein from cells without nicotinamide administration (-NAM), with nicotinamide (+NAM), PJ34 (PJ) administration, or sorted into GP2+ and GP2− populations. Cells were also treated with the sirtuin inhibitor EX527 or the sirtuin activator SRT1720. C, Quantification of the optical density of the 90 kDa line, average of 3 biological replicates. * indicates p < 0.05 in a paired Students t-test.

### Nicotinamide and differentiation affects mitochondrial oxidation in human pancreatic endocrine progenitors

Changes in the NAD^+^/NADH ratio (Fig. 2), in sirtuin activity (Fig. 3), and the lack of response to rotenone and FCCP in two-photon microscopy (Suppl. Fig. 1) suggested that mitochondrial activity is altered both over the course of differentiation and in the presence or absence of nicotinamide. To quantify mitochondrial function, we measured oxidative and glycolytic metabolism using the Seahorse metabolic profiler (Fig. 4A) of differentiating pancreatic progenitors after nicotinamide treatment for 2 or 5 days in stage 4 of the differentiation protocol. We hypothesized that as these cells differentiated towards pancreatic endocrine progenitors expressing Nkx6.1, that their metabolic activity would increase, and that nicotinamide exposure would also alter metabolism. Consistent with this hypothesis, the basal oxygen consumption rate (OCR) increases significantly from 2 days to 5 days of nicotinamide treatment (Fig. 4B). At day 5 nicotinamide appears to decrease the basal OCR, but is only significant to a 94% confidence level this timepoint (Fig. 4B). The difference in OCR becomes more apparent at day 5 with measurement of the maximum OCR after exposure of the cells to FCCP, in which cells treated with nicotinamide have lower maximum OCR than those not exposed to nicotinamide, and those treated with PARP inhibitor PJ34 (Fig. 4C). This would be consistent with active sirtuin inhibition of nicotinamide given that one of the main functions of the sirtuin family is to increase mitochondrial oxidation capabilities and rate. Extracellular acidification rate (ECAR), a measurement of glycolytic activity, showed decreased rates in day 2 of nicotinamide compared to day 5 (Fig. 4D). Similar to the measurements of OCR maximum, at day 5 cells treated with nicotinamide had significantly increased ECAR compared to untreated cells or those treated with PJ34 (Fig. 4D). Across all measurements, GP2+ and GP2− populations showed no metabolic differences, suggesting that an Nkx6.1 enriched population did not have significantly different metabolic activity. Palmitate addition, which usually increases oxygen consumption in metabolically mature cells did not change oxidative metabolism in differentiating hPSC and trended towards a decreased oxygen consumption rate, showing a defect in fatty acid metabolism at both day 10 and day 13 stages of differentiation (Suppl. Fig. 3).

**Figure 4:**
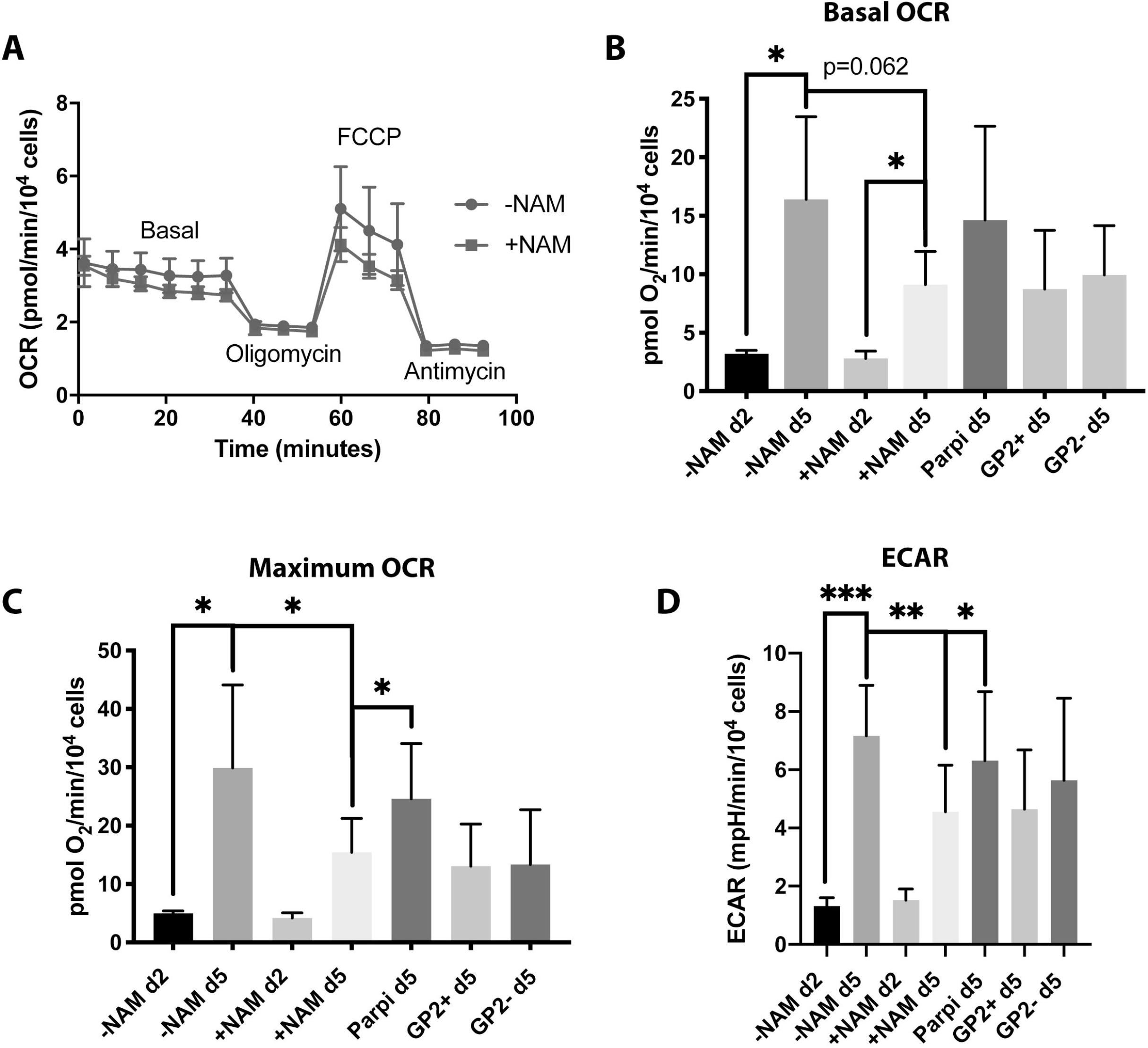
Metabolic rate increases as differentiation to pancreatic progenitors progresses and is insensitive to nicotinamide. A, Basal oxygen consumption rate (OCR) at day 2 and day 5 of nicotinamide or PJ34 (Parpi) treatment. Cells were also sorted into GP2+ and GP2− populations before measuring OCR. B, Maximum oxygen consumption rate at day 2 and day 5 in the presence or absence of nicotinamide, Parpi, and sorted into GP2+ and GP2− populations. C, Representative tracing of oxygen consumption rate at day 2 in the presence or absence of nicotinamide (NAM) when exposed to oligomycin, FCCP, and antimycin. D, Extracellular acidification rate (ECAR) at day 2 and day 5 in the presence or absence of nicotinamide, Parpi, and sorted into GP2+ and GP2− populations. * indicates p < 0.05, ** indicates p < 0.01, *** indicates p < 0.001, in a paired Students t-test.

### Isolated mouse islets exposed to nicotinamide increase mRNA transcripts related to beta cell identity

In order to determine if nicotinamide’s effects on human pancreatic progenitors were similar in other species, we isolated islets from 8 week old mice and exposed them to nicotinamide for 48 hours. We performed qRT-PCR for multiple transcripts including Nkx6.1, Pdx1, Ins1, Ins2, all essential genes for beta cell identity, as well as Nrf4a2, a downstream target of Nkx6.1, and DNMT1 which has no known function in the beta cell transcription network (Fig. 5). Nkx6.1, Pdx1, Ins2, and Nr4a2 were all significantly increased by exposure to nicotinamide, while DNMT1 was unchanged and Ins1 increased with a p-value of 0.0513. Exposure of mouse islets in vitro to nicotinamide increases mRNA transcripts related to beta cell function. Nicotinamide affects Nkx6.1 expression in both hPSC and mouse islets in a similar fashion, implying that there is a fundamental similarity in their regulation by PARP family inhibition.

**Figure 5:**
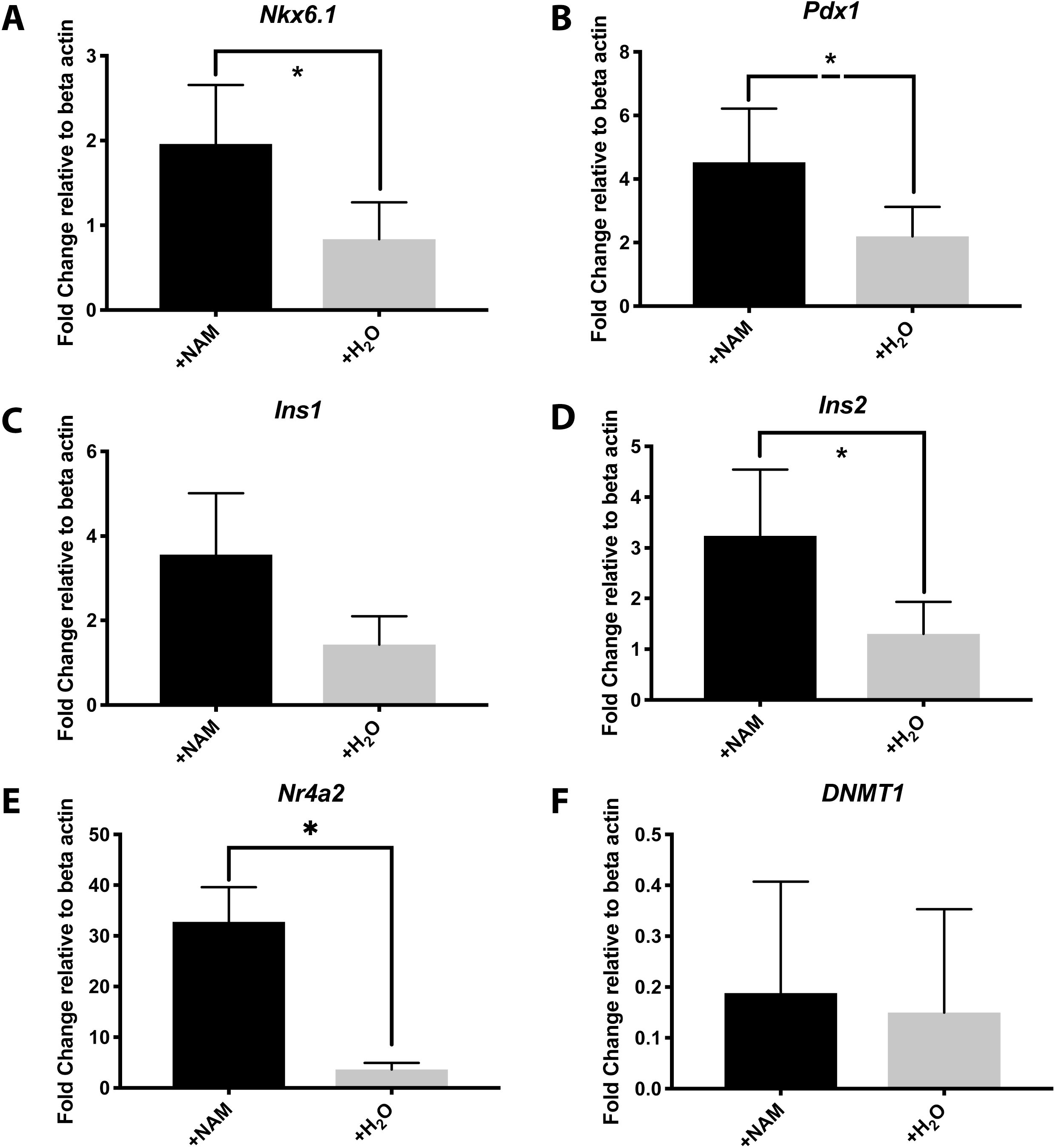
Fold change of transcripts in isolated mouse islets after 48 hours of nicotinamide or water treatment. A, Nkx6.1. B, Pdx1. C, Insulin 1. D, Insulin 2. E, Nr4a2. F, DNMT1. * indicates p < 0.05 in a paired Students t-test.

## Discussion

Nkx6.1 is an essential beta cell transcription factor whose effects have been well-characterized, but the upstream control of its expression has been unclear and predominately attributed to the transcription network involving other essential beta cell transcription factors. Nkx6.1 downregulation is an early marker of beta-cell dysfunction [27], and also has the largest downregulation of important beta cell transcripts in response to 10 weeks of mild hyperglycemia [28]. An understanding of the factors controlling its expression may be beneficial for developing ways of preserving or rescuing beta cell function in type 2 diabetes.

The finding that PARP inhibition is necessary for Nkx6.1 expression in hPSC suggests that Nkx6.1 responds to either a) cellular stress, expressed through the activity of the PARP family, which must be inhibited to allow Nkx6.1 expression, or b) responds to changes in the NAD^+^/NADH ratio, reflective of the redox state of the cell, which increases when PARP activity is reduced. The role of Nkx6.1 as an energy/redox sensing transcription factor is supported by the evidence that its knockout in mouse beta cells decreases the rate of glycolysis due to loss of Glut2 expression and reduces the amount of ATP found in the beta cell [16].

As hPSC differentiate they undergo a switch from glycolysis dependent to oxidative phosphorylation dependent metabolism [23], resulting in an increased capacity for ATP production accompanied by increased oxygen consumption rates. Consistent with this observation, we showed that cells increase their oxygen consumption rate as they transition from Nkx6.1 negative pancreatic progenitors on day 2 of nicotinamide treatment to Nkx6.1 positive endocrine progenitors on day 5 of nicotinamide treatment. In a similar but converse finding, Nkx6.1 expression decline was correlated with mitochondrial DNA reduction in diabetic Goto Kakizaki rats [29].

The sorting of cells into an Nkx6.1-enriched population expressing GP2 showed that the metabolic state of these human pancreatic endocrine progenitors remained similar at baseline regardless of the degree of Nkx6.1 expression. This was unexpected given the association of Nkx6.1 with increased ATP levels [16]. The GP2+ and GP2− populations were sorted from an original population that had been exposed to nicotinamide, and the basal metabolic rates are consistent with the unsorted population treated with nicotinamide. In contrast, the population treated with PARP inhibitor has a higher basal metabolic rate more consistent with the condition where nicotinamide was absent. These results suggest that any metabolic change triggered by Nkx6.1 occurs at a later point in differentiation, or that the inhibitory effects on metabolic rates caused by nicotinamide continued to affect both GP2+ and GP2− populations. It also shows that the observed metabolic changes are not the driver behind differentiation to an Nkx6.1+ population, but are instead an artifact of sirtuin inhibition by nicotinamide. The fact that the differentiated hPSC do not respond to palmitate further suggests that pancreatic progenitors and endocrine progenitors are metabolically immature cells, unable to efficiently convert fatty acids into energy.

In much earlier investigations, nicotinamide was found to rescue streptozotocin-induced beta cell death [14]. The results from this study show that nicotinamide support the preservation of pancreatic beta cell transcription factor expression in mouse islets, including *Pdx1, Nkx6.1* and *Ins2*. Nicotinamide exposure was also associated with upregulation of *Nr4a2. Nr4a1, Nr4a2, and Nr4a3,* which are orphan nuclear receptors that are bound to and regulated by Nkx6.1 [16]. Furthermore, the overexpression of Nkx6.1 stimulates beta cell proliferation through Nr4a1 and Nr4a3 induction [30], Nr4a3 genetic variants are associated with increased insulin secretion in humans [31], Nr4a receptors in conjunction with retinoid X receptors have been shown to upregulate carnitine palmitoyltransferase 1A (CPT1A)[32], and also respond to lipotoxicity in murine pancreatic beta cells [33]. Together, these findings link Nkx6.1 expression to the metabolic state of the pancreatic beta cell.

Nicotinamide exposure in this work is essential for robust Nkx6.1 expression in hPSC differentiating to pancreatic endocrine progenitors, and it acts predominately through PARP inhibition. While nicotinamide inhibits the sirtuin family and alters oxidative metabolic rates, these changes are correlated with but not necessary for Nkx6.1 expression. There are also global changes to metabolic oxidation as hPSC undergo differentiation, with oxidative rates increasing as differentiation progresses. Selective targeting of the NAD^+^ levels in adult beta cells may be a promising avenue for diabetes therapy.

## Acknowledgements

Thank you to Cynthia Fisher and Laura Prochazka for editing this manuscript. This work has been funded by the CIHR Foundation grant 375257 to P.W.Z., the Juvenile Diabetes Research Fund (JDRF), Beta Cell Biology Consortium (BCBC). C.W. is the recipient of CIHR Alexander Graham Bell Canadian Graduate Scholarship. P.W.Z. is the Canada Research Chair in Stem Cell Bioengineering.

---

**Supplementary Figure 1:**
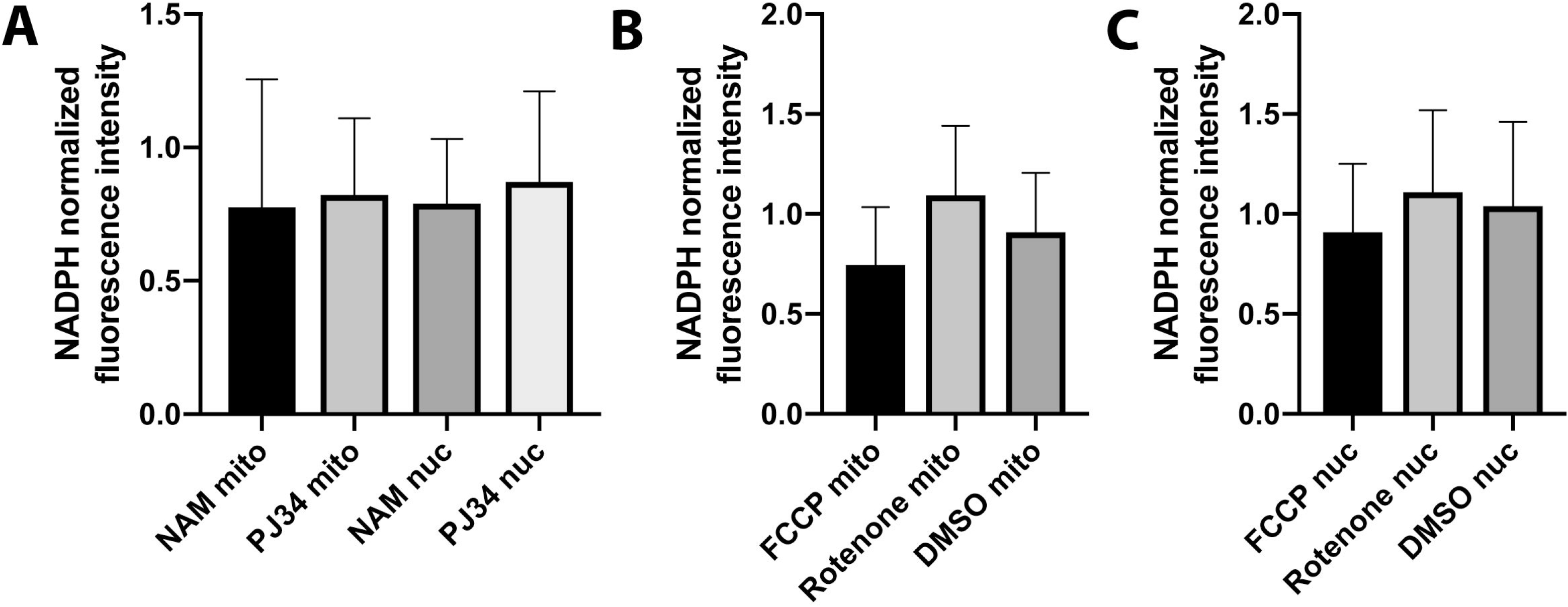
A, NADPH levels measured using 2-photon microscopy are unchanged in the mitochondria (mito) and nucleus (nuc) despite nicotinamide (NAM) and PJ34 administration. B, NADPH levels in the mitochondria are not significantly altered by FCCP, rotenone, or DMSO administration. C, NADPH levels in the nucleus are not significantly altered by FCCP, rotenone, or DMSO administration.

**Supplementary Figure 2:**
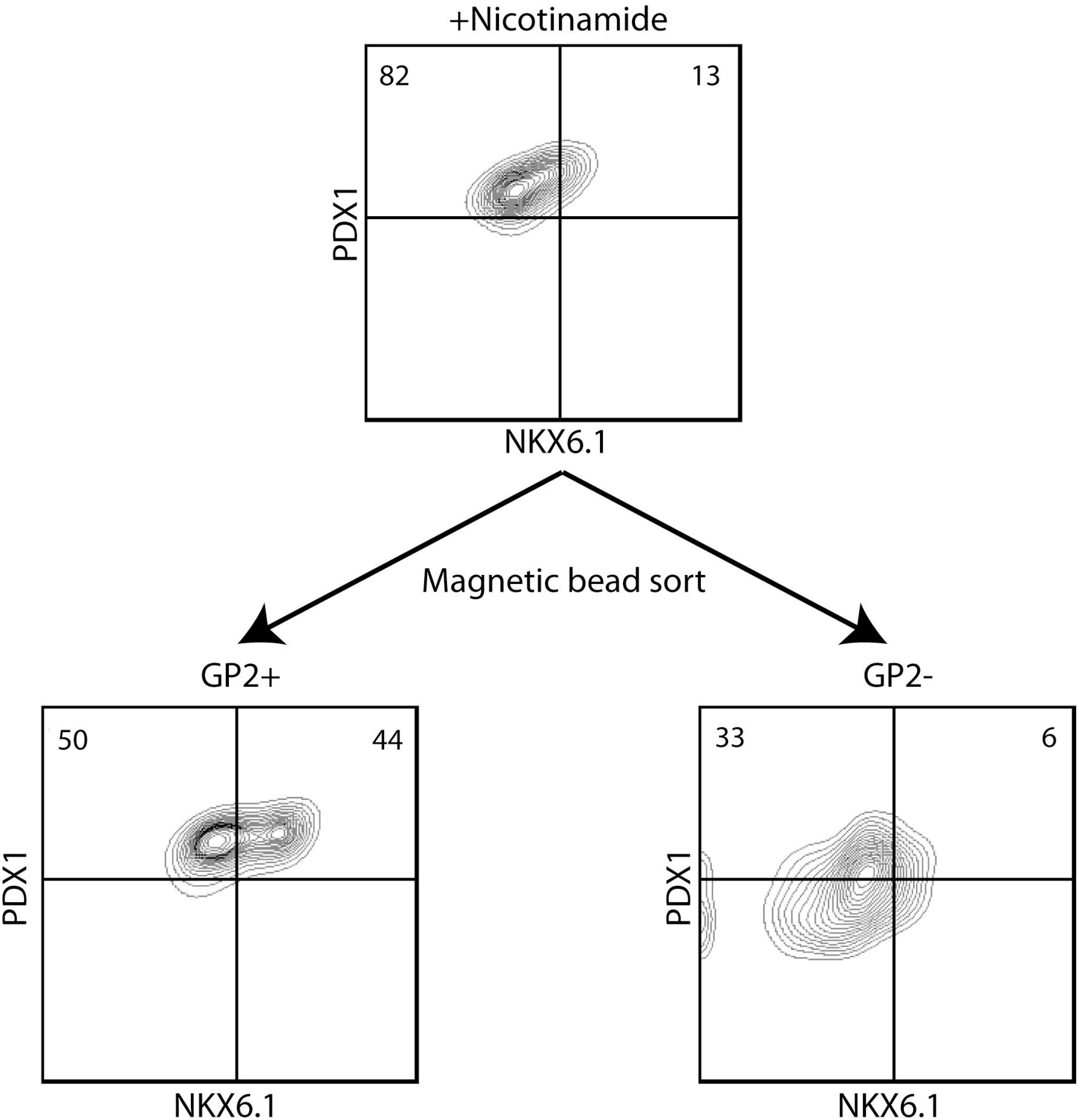
Representative example of magnetic bead sorting of a nicotinamide treated day 13 pancreatic progenitor population into GP2+, Nkx6.1 enriched and GP2-, Nkx6.1 deplete populations that were used for the experiments shown in figures 3–5.

**Supplementary Figure 3:**
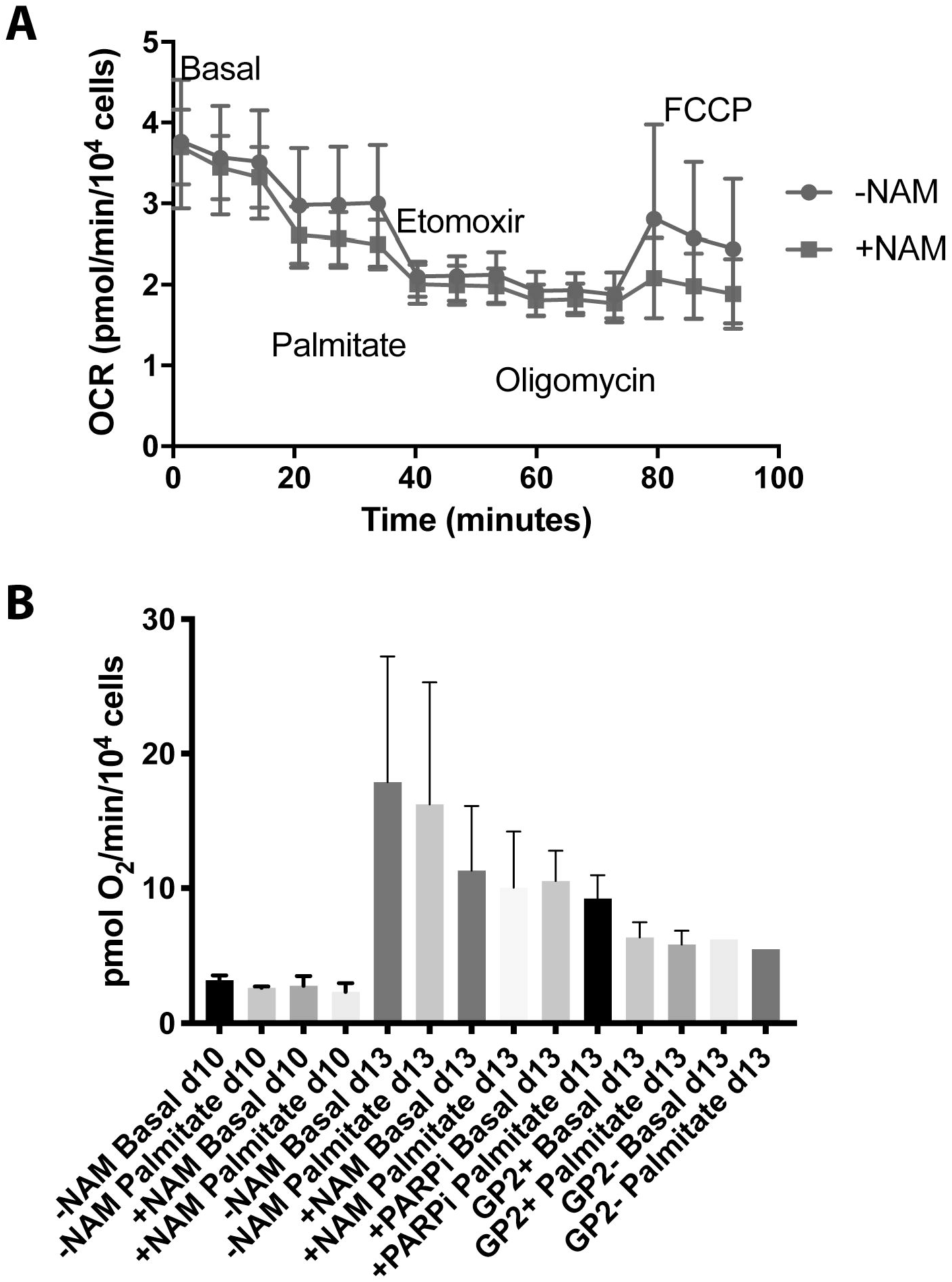
A, Representative oxygen consumption rate at day 2 of nicotinamide (NAM) treatment in the presence of palmitate, etomoxir, oligomycin, and FCCP in the presence or absence of NAM. B, Average oxygen consumption rates in the presence of palmitate, NAM, Parpi at day 2 and 5 of nicotinamide treatment and sorted into GP2+ and GP2− populations.

